# Tagless LysoIP method for molecular profiling of lysosomal content in clinical samples

**DOI:** 10.1101/2024.05.17.594681

**Authors:** Daniel Saarela, Pawel Lis, Sara Gomes, Raja S. Nirujogi, Wentao Dong, Eshaan Rawat, Sophie Glendinning, Karolina Zeneviciute, Enrico Bagnoli, Rotimi Fasimoye, Cindy Lin, Kwamina Nyame, Fanni A. Boros, Friederike Zunke, Frederic Lamoliatte, Sadik Elshani, Matthew Jaconelli, Judith J. M. Jans, Margriet A. Huisman, Christian Posern, Lena M. Westermann, Angela Schulz, Peter M. van Hasselt, Dario R. Alessi, Monther Abu-Remaileh, Esther M. Sammler

**Affiliations:** Medical Research Council (MRC) Protein Phosphorylation and Ubiquitylation Unit, School of Life Sciences, University of Dundee; Dow Street, Dundee DD1 5EH, U.K; Aligning Science Across Parkinson’s (ASAP) Collaborative Research Network; Chevy Chase, MD, 20815, USA; Department of Chemical Engineering, Stanford University; Stanford, CA 94305, USA; Department of Genetics, Stanford University; Stanford, CA 94305, USA; The Institute for Chemistry, Engineering and Medicine for Human Health (Sarafan ChEM-H), Stanford University; Stanford, CA 94305, USA; Department of Molecular Neurology, University Hospital Erlangen, Friedrich-Alexander-University Erlangen-Nürnberg (FAU); 91054, Erlangen, Germany; The Phil & Penny Knight Initiative for Brain Resilience at the Wu Tsai Neurosciences Institute, Stanford University; Stanford, CA 94305, USA; Department of Metabolic diseases, Wilhelmina Children’s Hospital, University Medical Center Utrecht, University Utrecht; Utrecht, The Netherlands; Department of Pediatrics, University Medical Center Hamburg-Eppendorf, Hamburg, Germany; Molecular and Clinical Medicine, Ninewells Hospital and Medical School, University of Dundee; Dundee, UK

## Abstract

Lysosomes are implicated in a wide spectrum of human diseases including monogenic lysosomal storage disorders (LSDs), age-associated neurodegeneration and cancer. Profiling lysosomal content using tag-based lysosomal immunoprecipitation (LysoTagIP) in cell and animal models allowed major discoveries in the field, however studying lysosomal dysfunction in human patients remains challenging. Here, we report the development of the “tagless LysoIP method” to enable rapid enrichment of lysosomes, via immunoprecipitation, using the endogenous integral lysosomal membrane protein TMEM192, directly from clinical samples and human cell lines (e.g. induced Pluripotent Stem Cell (iPSCs) derived neurons). Isolated lysosomes are intact and suitable for subsequent multimodal omics analyses. To validate our approach, we employed the tagless LysoIP to enrich lysosomes from peripheral blood mononuclear cells (PBMCs) derived from fresh blood from patients with CLN3 disease, a neurodegenerative LSD. Metabolic profiling of isolated lysosomes showed massive accumulation of glycerophosphodiesters (GPDs) in patients’ lysosomes. Interestingly, a patient with a milder phenotype and genotype displayed lower accumulation of lysosomal GPDs, consistent with their potential role as disease biomarkers. Altogether, the tagless LysoIP provides a framework to study native lysosomes from patient samples, identify novel biomarkers and discover human-relevant disease mechanisms.

## Introduction

Lysosomes are membrane-bound organelles whose function is critical to maintain cellular homeostasis through nutrient recycling, clearance of cellular waste and signaling (1-3).

Dysfunctional lysosomes are implicated in a wide range of human diseases, in particular lysosomal storage disorders (LSDs) and neurodegeneration (3-5). Lysosomes make up only 1-3% of a cell’s volume (6), which makes it challenging to interrogate changes in their molecular content under disease conditions. Recently, the LysoTagIP approach has enabled rapid immunoprecipitation of intact lysosomes, allowing subsequent analyses of their content and leading to a better understanding of human disease (6, 7). This approach is based on exogenous expression of the lysosome-resident transmembrane TMEM192 protein containing three tandem HA epitopes at its C-terminus (LysoTag) in target cells and mouse models, allowing for rapid lysosome immunoprecipitation.

Tagging patient lysosomes is impossible, thus, for studying lysosomal biology and dysfunction in clinical samples, we developed the “tagless LysoIP” method that enables the enrichment of lysosomes from a wide range of human-derived cells and clinical samples. The tagless LysoIP approach exploits an antibody capable of recognizing endogenous TMEM192, thereby bypassing the need to exogenously express a LysoTag. Using proteomic, metabolomic and lipidomic mass-spectrometry as well as flow cytometry and enzyme assays, we demonstrate that this method efficiently enriches lysosomes from human cell lines, human peripheral blood mononuclear cells (PBMCs), and human iPSC-derived neurons.

To further validate the utility of our method to study human disease, we applied the tagless LysoIP to profile the metabolite content of CLN3 patients’ lysosomes. CLN3 disease is the most common form of neuronal ceroid lipofuscinoses (NCLs), a neurodegenerative LSD family with childhood onset, progressive degeneration of the brain, including retina and intracellular accumulation of autofluorescent lipopigment (lipofuscin) (8). Recent work using LysoTag mice and cells revealed that the CLN3 protein is required for the clearance of glycerophosphodiester (GPDs) from lysosomes and that GPDs accumulate in CLN3 knockout mice and cells harboring the LysoTag (9, 10). Here we show significant accumulation of GPDs in lysosomes derived from patients’ PBMCs using the tagless LysoIP method compared to those from healthy controls; a finding that would have gone unnoticed at the whole cell lysate level.

Our results indicate that the tagless LysoIP method is a powerful and rapid approach to enrich lysosomes from clinical samples and other human cells such as iPSC-derived neurons for both biomarker discovery and understanding disease pathology.

## RESULTS

### Development of the tagless LysoIP method

TMEM192 is a 271-residue lysosomal protein, consisting of 4 transmembrane domains, with its N- and C-termini facing the cytoplasm (fig. S1A). We identified two rabbit monoclonal antibodies, TMEM192^AB1^ (Abcam ab186737) and TMEM192^AB2^ (Abcam ab185545) that successfully immunoprecipitate the human TMEM192-3HA protein and detect its presence with immunoblotting in lysates of HEK293 cells stably expressing the protein (fig. S1B and S1C). Epitope analysis revealed that TMEM192^AB1^ recognizes C-terminal residues between 235-250, while TMEM192^AB2^ recognizes those between 200-235 (fig. S1C). To test for the ability of the identified antibodies to enrich for lysosomes from detergent free homogenized cell lysates, we covalently coupled them to magnetic beads and incubated each conjugated antibody with homogenates from HEK293 cells expressing the LysoTag (TMEM192-3HA). As controls, we used magnetic beads coupled to HA antibodies (HA-IP) as well as bovine serum albumin (BSA/MockIP) (Fig. 1A). Cells were homogenized using a ball bearing homogenizer in an isotonic potassium phosphate-buffered saline (KPBS) with protease and phosphatase inhibitors (Fig. 1B). To minimize leakage, the homogenization, immunoprecipitation and wash steps were optimized to take ∼10 minutes as in our original LysoTagIP protocol (Fig. 1B) (11).

**Fig. 1.**
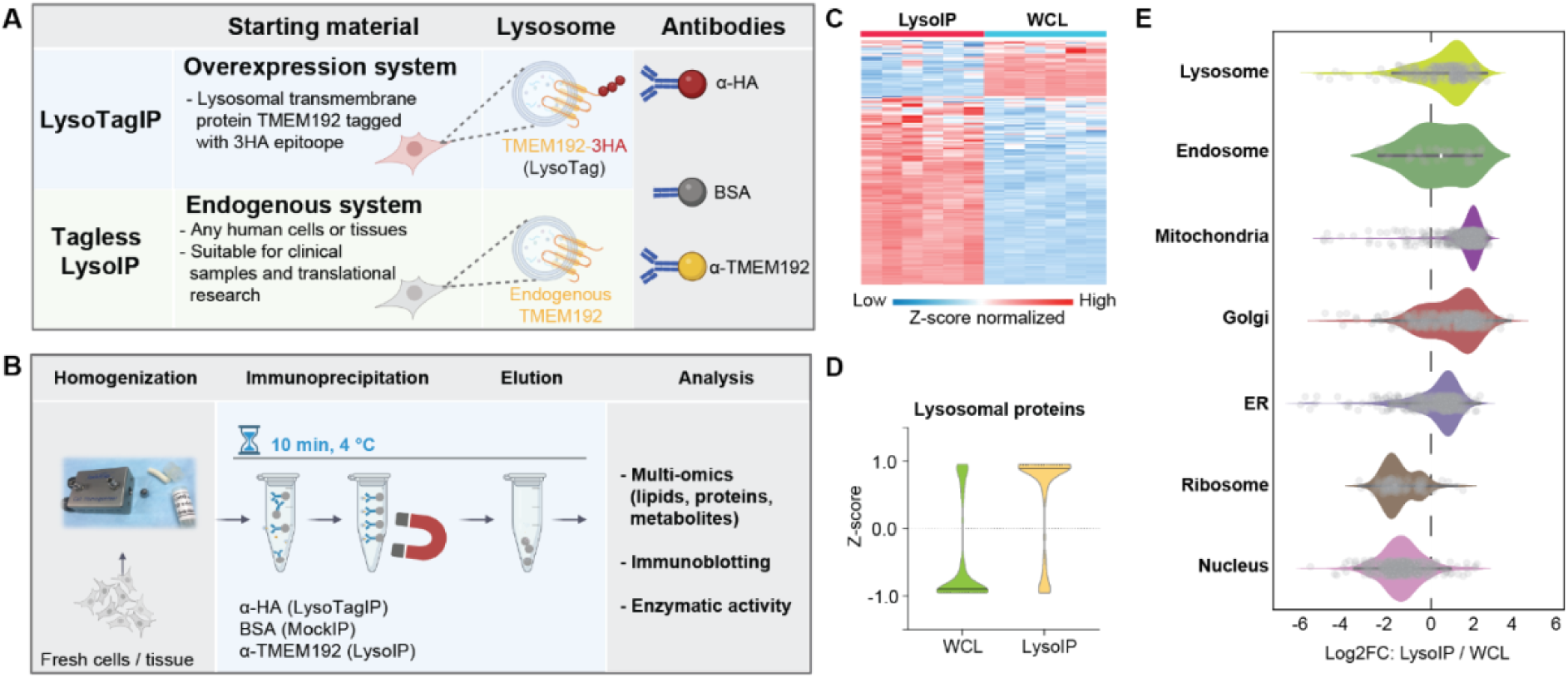
Tagless LysoIP for enriching lysosomes from clinical samples. **(A)** Concept of the tagless LysoIP method via immunoprecipitation of the lysosomal transmembrane TMEM192 protein in comparison to the LysoTag system that relies on overexpression of 3HA epitopes at the TMEM192 C-terminus. **(B)** Tagless LysoIP workflow, **(C)** Protein profile heatmap. **(D)** Violin plots of lysosomal proteins enriched via the tagless LysoIP in the immunoprecipitates and whole cell lysates (WCLs) from wildtype HEK293 cells. **(E)** Organelle profiling demonstrates significant enrichment of lysosomal but also non-lysosomal proteins.

To rigorously assess the enrichment of lysosomal proteins in immunoprecipitates (IPs), whole cell lysates (WCLs) and IPs were initially analyzed using high-resolution data-independent acquisition (DIA) liquid chromatography/tandem mass spectrometry (LC/MS-MS). TMEM192^AB1^ and TMEM192^AB2^ IPs clustered closely together in both wildtype and LysoTag expressing cells (fig. S2A). As expected, the HA-IP from LysoTag cells yielded robust enrichment of lysosomal proteins (fig. S2B). TMEM192^AB1^ also significantly enriched lysosomes in wildtype and LysoTag cells, albeit to a lower extent than observed in HA-IPs from LysoTag cells (fig. S2B and 2C). TMEM192^AB2^ antibody was less efficient at enriching lysosomes than TMEM192^AB1^ antibody (fig. S2B and S2C). Thus, all subsequent work was performed using TMEM192^AB1^ antibody, and the approach was termed the “tagless LysoIP method”.

Further proteome analyses established that lysosome-annotated proteins are enriched in TMEM192^AB1^ IP from wildtype HEK293 cells (Fig. 1D and 1E, fig. S2B to S2D). These include key lysosomal proteins such as LAMP1, LAMTOR1, TMEM55B, CTSC, CTSD and GBA1 (fig. S2E).

Despite the clear enrichment of lysosomal content, organelle profiling revealed the presence of non-lysosomal proteins in the tagless LysoIP, especially from endosomal, mitochondrial and Golgi compartments (Fig. 1E). This was further illustrated in the Volcano plot analysis (fig. S2D) and when looking at the relative intensities of representative cytoplasmic (Tubulin) and other organelle proteins (VDAC1; mitochondria, ACBD3; Golgi) in LysoIP compared to MockIP and WCL samples (fig. S2E). Altogether, these data suggest that the tagless LysoIP enriches for lysosomes while minor fractions from other compartments are also present.

To test that the presence of mitochondrial proteins in the LysoIPs was not due to nonspecific binding of the TMEM192^AB1^ antibody to mitochondria, we co-stained wildtype HEK293 cells with the TMEM192^AB1^ antibody and either a lysosomal (LAMP1) or mitochondrial marker (ATPB) (fig. S3A). This showed that binding of the antibody is lysosome specific, suggesting that the presence of mitochondrial proteins in the LysoIP is likely due to other factors than mere binding of the TMEM192^AB1^ antibody to mitochondria.

To confirm that the enriched lysosomes are intact, we preincubated HEK293 cells with the pH-dependent lysosomal fluorescent dye LysoTracker prior to cell homogenization (fig. S3B). IPs were subsequently analyzed by flow cytometry, which demonstrated a marked increase of LysoTracker labeled lysosomes in LysoIPs compared to MockIPs, indicating small molecule retention during the tagless LysoIP process (fig. S3C and S3D). Pretreatment of HEK293 cells with bafilomycin to suppress the acidification of lysosomes reduced LysoTracker signal by ∼50%. Moreover, the activity of cathepsin D, a luminal lysosomal enzyme, was significantly higher in LysoIP compared to MockIP and WCL, further indicating that the tagless LysoIP enriches intact lysosomes (fig. S3E).

We next performed targeted lipidomic analysis for bis(monoacylglycero)phosphates (BMPs) as these are relevant biomarkers for a range of neurodegenerative conditions. For example, BMPs are elevated in urine of patients with LSDs (11) and may have utility as a urine biomarker in leucine-rich repeat kinase 2 (LRRK2) associated Parkinson’s disease (PD) (13 - 15). We were able to robustly measure BMPs in HEK293 cells and found that they were enriched in LysoIP compared to MockIP samples (fig. S4).

To test the tagless LysoIP method in relevant cellular models of human disease, we performed proteome analysis on lysosomes enriched from iPSC-derived dopaminergic neurons (fig. S5). Principal component analysis revealed that LysoIP samples clustered distinctly from their corresponding WCLs while heatmap clustering revealed clear enrichment of lysosomal proteins in the LysoIPs compared to WCLs (fig. S5C and S5D). This was also seen in the volcano plot analysis comparing LysoIP to whole cell proteomic content (fig. S5E), which again highlighted the presence of mitochondrial proteins in the LysoIPs. As with the HEK293 experiments, proteins from other organelles were also present, although to a lower extent than the lysosomal ones (fig. S5F); the lysosomal proteins LAMP1, LAMTOR1, CLN3, CTSD, and beta-hexosaminidase A were enriched in LysoIPs compared to WCLs (fig. S5H), while most neuronal markers (bIII Tubulin, alpha-synuclein, and the dopamine metabolism-associated proteins tyrosine hydroxylase (TH) and ALDH1A1) were depleted in the LysoIP compared to WCL (fig. S5H). Cytoplasmic markers including GAPDH, actin and tubulin were depleted while the Golgi-associated ABCD3 protein was enriched in LysoIPs (fig. S5I). Interestingly, the mitochondrial VDAC1 protein was not enriched in LysoIPs (fig. S5I).

### The tagless LysoIP method enriches lysosomes from PBMCs

Human peripheral mononuclear cells (PBMCs) are routinely isolated from human peripheral blood for clinical and biomarker studies. These comprise a heterogeneous group of white blood cells including lymphocytes (T cells, B cells, and NK cells), monocytes, and dendritic cells (16). We explored the feasibility of performing lysosomal enrichment from PBMCs isolated from fresh whole blood via the tagless LysoIP method as depicted in Fig. 2A. We first undertook pilot experiments with and without the potent elastase protease inhibitor, diisopropylfluorophosphate (DIFP) in the cell homogenization buffer as PBMCs are often contaminated with trace amounts of neutrophils that contain high levels of elastases that can degrade proteins in extracts (17). The addition of DIFP in the homogenization buffer significantly increased the detectability of key organelle marker proteins (LAMP1, TMEM55B, LAMTOR1, GM130, HSP60), not only in the IPs but also in WCLs. Cytoplasmic proteins such as tubulin and GAPDH on the other hand were significantly depleted in LysoIPs compared to WCLs (fig. S6). This emphasizes the importance of maintaining DIFP in the homogenization buffer for all PBMC tagless LysoIP experiments. It should be noted that DIFP is a highly toxic organophosphate and thus must therefore be handled with appropriate care.

**Fig. 2.**
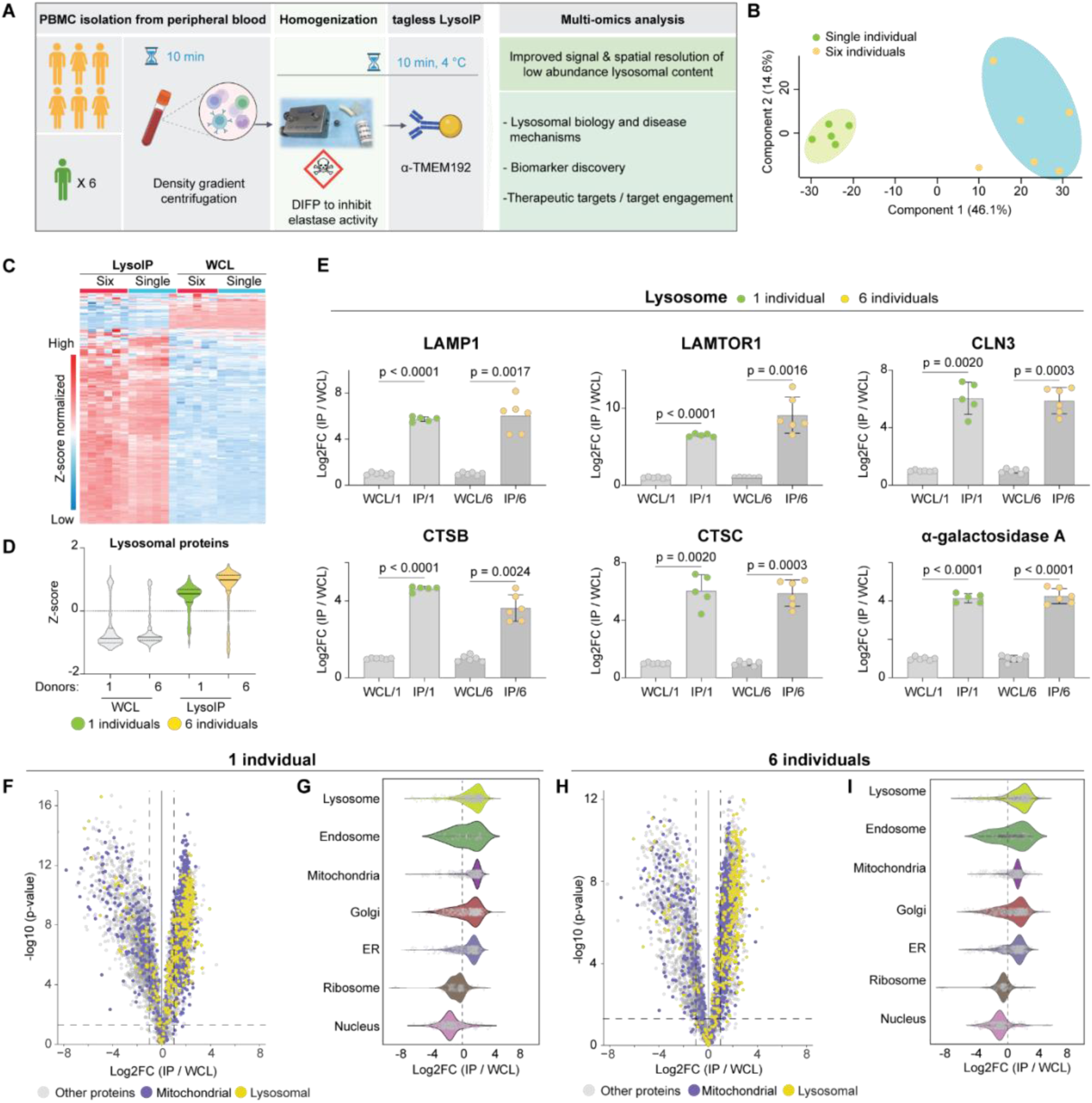
Tagless LysoIP in human peripheral blood mononuclear cells (PBMCs). **(A)** Workflow **(B)** Principal component analysis of DIA mass-spectrometry data of tagless LysoIPs from PBMCs from a single donor (6 replicates) to test technical variability and 6 different donors to test biological variability **(C)** Protein profile heatmap. **(D)** Lysosomal protein enrichment. **(E)** Bar plots of representative lysosomal transmembrane (top panel) and intraluminal (lower panel) proteins in LysoIPs from 1 and 6 donors compared to the respective whole cell extracts. Volcano and violin plots of LysoIPs compared to whole cell extracts and organelle profiling from the single **(F, G)** and multiple **(H, I)** donor experiments. Curtain links: https://curtain.proteo.info/#/b573afb8-df5d-4a39-b5ba-88eb9488820d **(F)**, https://curtain.proteo.info/#/83006f89-901d-44db-b078-5b8504844ee6 **(H)**.

To test technical reproducibility and inter-individual variability of the tagless LysoIP methodology in PBMCs, we performed 6 tagless LysoIPs from a single healthy donor (technical replicates) as well as tagless LysoIPs from 6 healthy individuals (biological replicates) (Fig. 2A, fig. S7). The samples were analyzed in parallel by DIA LC/MS-MS as before. On average, over 5000 unique proteins were identified in the LysoIP samples, except for 1 replicate from the single donor experiment (1003 identified proteins) that was subsequently excluded from further analysis. Principal component analysis revealed that replicates from the single and multiple donor tagless LysoIP experiments clustered closely together, but as expected there was more variation between LysoIPs from multiple donors in the biological replicate experiment compared to the technical replicate experiment from a single donor (Fig. 2B). MockIPs were only performed for the technical variability single donor experiment, but also clustered closely but distinctly from the LysoIPs and WCLs (fig. S7A). Heatmap clustering of the DIA-MS data revealed that levels of lysosomal proteins were similar in all LysoIP samples (Fig. 2C). Likewise, violin plots (Z-score normalized) confirmed strong enrichment of lysosomal proteins in the tagless LysoIP experiments from human PBMCs compared to WCLs (Fig. 2D). Volcano plot analysis confirmed enrichment of lysosomal proteins in the LysoIP for the technical replicates (single donor) (Fig. 2F) and biological replicate (multiple donors) experiments (Fig. 2H), compared to corresponding WCLs. Levels of luminal (CTSB, CTSD, and GBA1) and transmembrane (LAMP1, LAMTOR1 and CLN3) lysosomal proteins displayed 4 to 10-fold enrichment in LysoIPs compared to whole cell extracts (Fig. 2E). We noted that Golgi (ACBD3 & GM130) (fig. S7B) and mitochondrial (HSP60, VDAC1) (fig. S7C) markers were also enriched 4 to 6-fold in PBMC LysoIPs compared to WCLs. Cytosolic markers such as Tubulin and Actin were depleted in PBMC LysoIP samples (fig. S7D). This may reflect the capture of the newly synthesized, tagless LysoIP antigen TMEM192 as it traverses the secretory pathway. Organelle profiling of PBMC LysoIPs for both single (Fig. 2G) and multiple donors (Fig. 2I) showed similar results to what had been observed in HEK293 cells (Fig. 1E) with regards to the presence of mitochondria and other organelles.

To confirm that the enriched lysosomes are intact, we preincubated PBMCs with the pH-dependent lysosomal fluorescent dye LysoTracker prior to cell homogenization and tagless LysoIP and subsequently analyzed the LysoIPs by flow cytometry (Fig. 3A). Consistent with isolation of intact lysosomes in the HEK293 cells (fig. S3), the tagless LysoIP enriched lysosomes from PBMCs that retained the LysoTracker signal, which was responsive to bafilomycin treatment (Fig. 3B, 3C). Furthermore, protein levels (Fig. 3D) and catalytic activity (Fig. 3E) of the intraluminal lysosomal hydrolase GCase (glucocerebrosidase, GBA1) were enriched in LysoIPs compared to WCL fractions. Of note, homozygous GBA1 variants cause Gaucher’s disease, a common LSD while heterozygous GBA1 variant carrier status increases risk for PD (18).

**Fig. 3.**
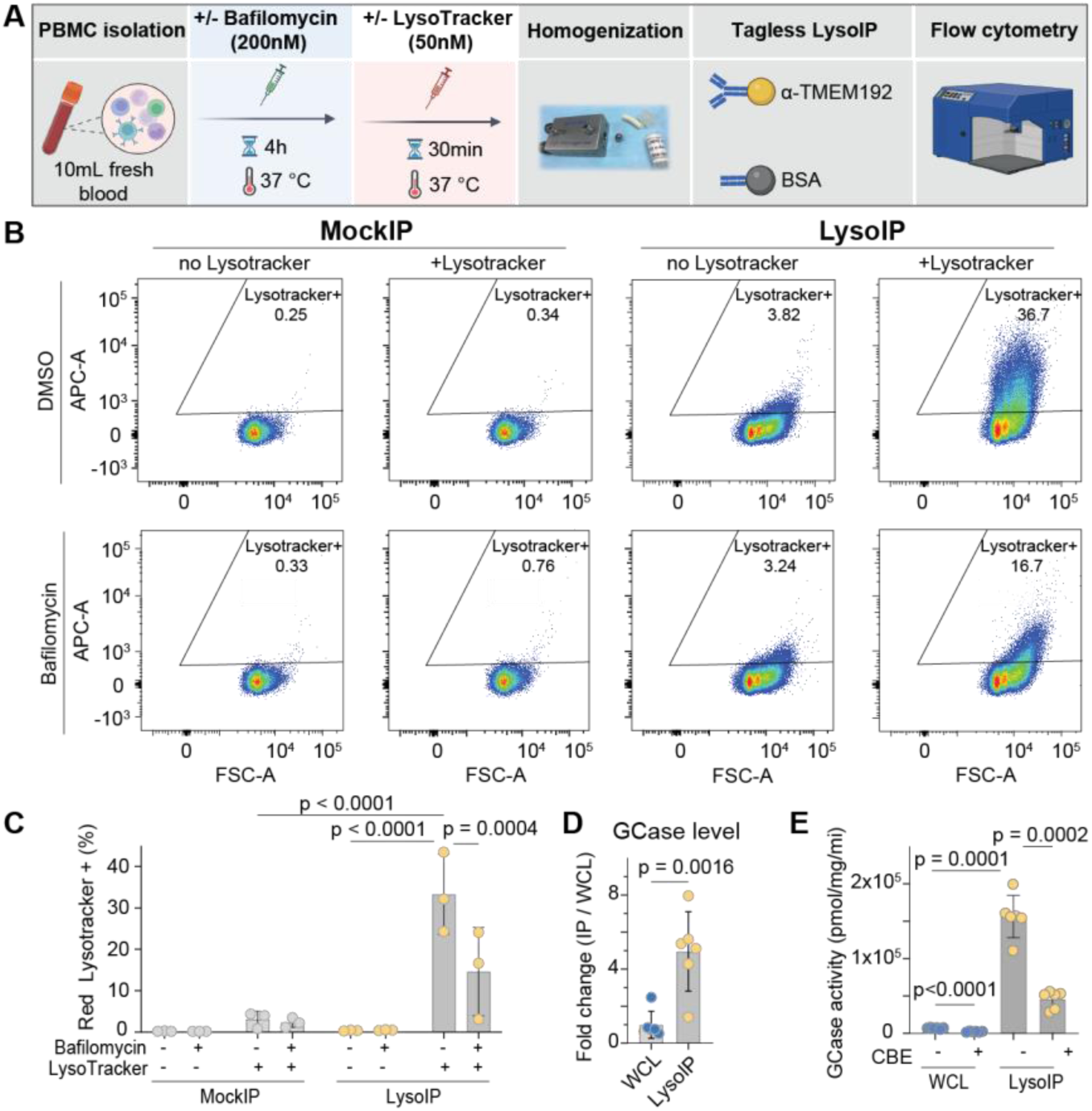
Tagless LysoIP enriches intact and functional lysosomes from healthy donor PBMCs. **(A)** Workflow of flow cytometry analysis of magnetic beads bound to lysosomes enriched via tagless LysoIPs or MockIPs from PBMC homogenates pretreated with and without the V-ATPase inhibitor bafilomycin A1 (200 nM), prior to staining with Red Lysotracker (50 nM). (**B)** Representative scatter plot from one of the three donors, (Y-axis) and bead size (Forward scatter, FSC, X-axis), **(C)** Quantification of the percentage of beads positive for Lysotracker fluorescence (N = 3 donors) and analysis by ordinary two-way ANOVA with Tukey’s multiple comparison test. **(D)** GCase (glucocerebrosidase, GBA1) protein enrichment quantified by DIA mass-spectrometry in LysoIP **(E)** GCase activity measured by 4-methylumbelliferone (4-MU) assay in lysosomes enriched from PBMCs from 6 donors.

### The application of tagless LysoIP in clinical settings

Encouraged by the ability of our method to enrich intact lysosomes from healthy donor PBMCs for multimodal profiling, we explored the feasibility and utility of the tagless LysoIP methodology to identify clinically relevant lysosomal biomarkers in LSDs.

We turned to CLN3-disease, a devastating early-onset neurodegenerative LSD, which we previously studied in human cell lines and animal models (9, 10). In total, fresh peripheral blood samples were collected from 10 individuals, 5 patients with CLN3-associated NCL disease and 5 sex-matched controls. 4 of the patients carried a common 1kb deletion mutation in the homozygous state and presented with typical juvenile onset with retinal disease and additional complex neurodegenerative symptoms. 1 patient possessed compound heterozygous mutations that may represent a partial loss of function and presented with adult onset (3rd decade) retinal only disease (Tab. 1).

**Table 1.** Overview of clinical study participants. Fresh peripheral blood for PBMC isolation and subsequent immunoaffinity enrichment of lysosomes via tagless LysoIP was collected from 5 patients with CLN3 disease and 5 controls. Four CLN3 patients carried the common 1kb deletion (*c.461-280_677+382del966) in the homozygous state and presented with classic juvenile CLN3 disease onset. One CLN3 patient (participant 10) presented with retinal-disease only with age at disease onset at 29. She carried bi-allelic compound heterozygous variants in CLN3 (**c.391_392del; p.(Ser131fs) and c.969G>A; r. (spl?)). Samples from participants 1/2/3, 4/5, 6/7 and 8/9/10 were collected at the same time. Age and age at disease onset in years (y) and months (m). Biological sex as female (F) and male (M), Collection sites: Hamburg (Ham) and Utrecht (Utr).

|  | Group | Genotype | Phenotype | Age | Age at onset | Sex | Site |
| --- | --- | --- | --- | --- | --- | --- | --- |
| 1 | CLN3 disease | hom 1kb deletion* | classic juvenile | 13y 2m | 4y 0m | F | Ham |
| 2 | Control | - |  | 24Y | - | M | Ham |
| 3 | Control | - |  | 24Y | - | F | Ham |
| 4 | CLN3 disease | hom 1kb deletion* | classic juvenile | 8y 10m | 6y 5m | F | Ham |
| 5 | Control | - |  | 27y | - | F | Ham |
| 6 | Control | - |  | 28y | - | F | Ham |
| 7 | CLN3 disease | hom 1kb deletion* | classic juvenile | 9y 1m | 5y 5m | M | Ham |
| 8 | Control | - |  | 26 | - | M | Utr |
| 9 | CLN3 disease | hom 1kb deletion* | classic juvenile | 8y | 5y | M | Utr |
| 10 | CLN3 disease | bi-allelic compound het variants** | retinal-only | 52y | 29y | F | Utr |

We first undertook untargeted metabolomic mass spectrometry to profile the metabolite content of enriched lysosomes (9) (Fig. 4A). This revealed marked elevation of 5 glycerophosphodiesters (GPDs) in patients’ lysosomes that we were able to unambiguously annotate, namely glycerophosphoglycerol (GPG), glycerophosphoinositol (GPI), glycerophosphocholine (GPC), glycerophosphoethanolamine (GPE) and glycerophosphospingosine (GPS) (Fig. 4A, B). Targeted analyses in both LysoIPs and corresponding whole cell fractions demonstrated massive accumulation of these metabolites in CLN3-disease lysosomes, with an average of 17 to 830-fold accumulation of all 5 annotated GPDs (GPG 830-fold, GPI 440-fold, GPS 95-fold, GPE 42-fold, GPC 17-fold) in the lysosomes from CLN3 patients with the classic 1 kb deletion compared to healthy controls (Fig. 4C). At the whole cell level, a significant increase was only observed for 1 GPD metabolite namely GPS (8-fold) whereas the remaining GPDs (GPG, GPI, GPC, GPE) did not show a significant increase despite showing a potential trend in some cases. These data are consistent with our findings in cell culture and mouse models (9, 10). Interestingly, the patient with the milder disease associated with compound heterozygous mutations in CLN3 still displayed an enrichment of GPDs in the LysoIP compared to its corresponding WCL fraction. However, this enrichment was to a lower extent of what was observed in patients with the classical, more severe genotype and disease manifestation, but still higher when compared to healthy controls (4 to 30-fold) (Fig. 4C). These results indicate that the tagless LysoIP can potentially be used to identify and monitor disease biomarkers in human LSDs.

**Fig. 4.**
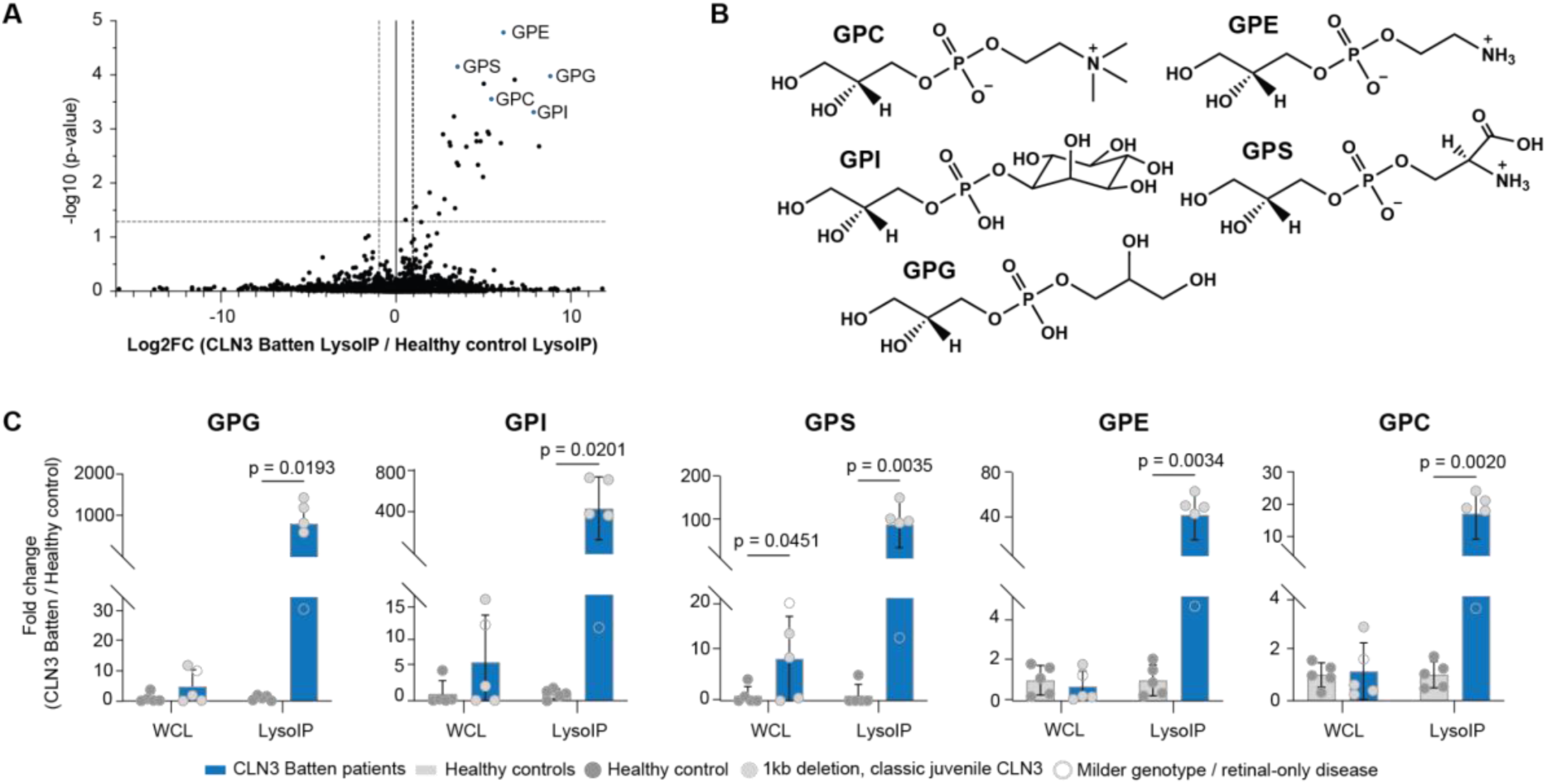
Striking GPD accumulation in lysosomes enriched from CLN3 disease patients. **(A)** Volcano plot comparing untargeted metabolomic LysoIP data derived from PBMCs from CLN3 patients and healthy controls. **(B)** The chemical structure of the annotated GPDs in this study. **(C)** Targeted analyses of GPDs in the LysoIPs and corresponding WCLs. Data presented as mean ± SEM (n = 5). Statistical analysis was performed using Student’s t-test.

## DISCUSSION

Lysosome dysfunction has been implicated in a myriad of human diseases. While tools to profile lysosomal content have been established for cell culture and animal models by expressing a tagged lysosomal membrane protein for organelle immunoprecipitation (9, 11), profiling lysosomal content from patients has remained a great challenge.

Here, we report the development of the “tagless LysoIP’’ method that allows the rapid enrichment of lysosomes via immunoprecipitation using an antibody against the integral lysosomal membrane protein TMEM192. After validating the method and confirming the intactness of the enriched lysosomes, a requirement to preserve their content, we applied the tagless LysoIP for the multimodal analyses of lysosomes from human cells including cultured cell lines, iPSC-derived neurons and human PBMCs. To test the utility of the method in uncovering molecular pathways involved in disease pathology at the subcellular level, we isolated and profiled lysosomes from PBMCs collected from CLN3-disease patients and healthy individuals. In striking agreement with our cell culture and mouse studies, we identified massive accumulation of GPDs in CLN3 patients’ lysosomes (9, 10). Of importance, the elevation of GPDs was barely observed and mostly not significant (except for GPS) at the whole cell level. These robust results emphasize the need of lysosomal enrichment via the tagless LysoIP method to study diseases where lysosomes are implicated including LSDs and neurodegenerative conditions, as most of the disease relevant alterations could otherwise be missed.

Interestingly, one of the CLN3 patients included in this study had a much delayed age of onset with two compound heterozygous mutations in *CLN3* causing an atypical mild phenotype restricted to retinal pathology only (Table 1). This individual still had elevated GPD metabolites within the lysosome, but at least 5-fold (between 4.8 to 43-fold) lower than in CLN3 patients that are homozygous for the common 1 kb deletion that results in a complete loss of function. This example illustrates the quantitative nature of the tagless LysoIP and its potential to measure lysosome-restricted biomarkers and correlate them to disease phenotypes, especially in cases with atypical mutations (19, 20).

While the tagless LysoIP represents a major step towards applying subcellular profiling to clinical samples, the LysoTag approach that relies on overexpressing the 3HA tagged TMEM192 remains the default option for other applications as it allows for the isolation of more content with higher purity (fig. S2B). We also found that the tagless LysoIP method is less efficient in enriching lysosomes from cryopreserved PBMCs compared to freshly isolated PBMCs. While our data indicate that TMEM192^AB1^ antibody does not co-localize to mitochondria using a representative marker (fig. S3A), it may still be that mitochondrial and other organellar proteins co-enriched in LysoIPs are a result of nonspecific contamination. Moreover, the TMEM192^AB1^ antibody is specific for the human protein and does not recognize the mouse TMEM192 protein due to amino acid sequence differences in the epitope. Thus, the current version of the tagless LysoIP is unsuitable for experiments in mouse models. Future work should aim at screening more antibodies that can resolve both issues and expand the utility of our optimized platform.

Altogether, our data show that the tagless LysoIP approach is a robust methodology for enriching lysosomes from clinical samples and human cell lines including human iPSC-derived neurons that provides a framework for studying lysosomal dysfunction in human diseases. The potential use of this novel technology spans from biomarker discovery, drug screening, to uncovering the function of lysosomal gene products. To determine the ultimate use as a biomarker to assess efficacy of experimental therapies, cross-sectional and longitudinal analyses of samples from a larger cohort of CLN3 patients are mandatory. In addition, these patients should be well characterized by established clinical scoring systems, e.g. Unified Batten Disease Rating Scale (22) or Hamburg CLN3 ophthalmic rating scale (23). Given that lysosomal dysfunction is a hallmark of cognitive decline in PD and other neurodegenerative disorders (24), the tagless LysoIP represents an important asset for translational research for studying lysosomes using clinical and iPSC-derived cell lines across a wide range of human diseases.

## Acknowledgments

We thank all the volunteers and patients for donating blood samples; without their generosity this study would not have been possible. We appreciate the support provided by the 2 clinical sites in Hamburg and Utrecht and thank Julia Vandrey (University Hospital Erlangen) for excellent support during the iPSC-differentiation. We acknowledge Dr. Herman van der Putten from the NCL-Stiftung (https://www.ncl-stiftung.de) for the helpful discussions and facilitating the clinical collaborations. We are grateful for the constructive feedback throughout the project provided by Suzanne Pfeffer (Stanford University) and Miratul Muqit (University of Dundee). We thank all our funders and acknowledge the excellent technical support from the MRC-Protein Phosphorylation and Ubiquitylation Unit (MRC PPU), in particular the mass-spectrometry team (Dr. Renata F. Soares) and MRC PPU Reagents and Services teams.

## Funding

This research received funding from the following sources:

Aligning Science Across Parkinson’s [ASAO-000463] through Michael J. Fox Foundation for Parkinson’s Research (MJFF) (DRA, MA-R)

UK Medical Research Council [grant number MC_UU_00018/1] (DRA and DRA lab members: DS, PL, RSN, RF, EB, MJ)

Pharmaceutical companies supporting the Division of Signal Transduction Therapy Unit [Boehringer Ingelheim, GlaxoSmithKline, and Merck KGaA] (DRA, ES)

Personal CSO Scottish Senior Clinical Academic Fellowship [SCAF/18/01] (ES)

Wellcome Trust PhD studentship [218520/Z/19/Z] (KZ)

Additional support was received from the NCL Foundations (NCL-Stiftung), Nächstenliebe e.V., German Center for Child and Adolescent Health, Beat Batten foundation for MA-R, the Netherlands and the Freemasons Berlin Outpost 46/ First Berlin Foundation

## Author contributions

Conceptualization: DRA, MAR, ES

Methodology: DS, PL, SG, RSN, WD, FL, KZ, ER, RF, EB, SG, FZ, FAB, MJ, MAH, DRA, MAR, ES

Investigation: DS, PL, SG, RSN, WD, KZ, FL, ER, CL, KN, RF, EB, SG, FZ, FAB, MJ, CP, LW, ES

Visualization: DS, PL, SG, RSN, WD, KZ, ER, RF, EB, MJ, ES

Funding acquisition: AS, PMvH, DRA, MAR, ES

Project administration: DRA, MAR, ES

Supervision: DRA, MAR, ES

Writing – original draft: DS, PL, RSN, DRA, ES

Writing – review & editing: DS, PL, SG, RSN, WD, KZ, ER, RF, CL, KN, EB, SG, FZ, FAB, MJ, JJMJ, MAH, CP, LW, AS, PMvH, DRA, MAR, ES

## Data and materials availability

All data needed to evaluate the conclusions in the paper are present in the manuscript and/or the Supplementary Materials. All the primary data presented here have been deposited in publicly accessible repositories. Immunoblotting data and confocal microscopy data (raw data files and their quantitation and statistical analysis) have been deposited in Zenodo (https://doi.org/10.5281/zenodo.11085342). Proteomic data have been deposited in the ProteomeXchange PRIDE repository (identifier: PXD052082). The dataset reviewer login details: Username. Password: Gd4QFVsL. All plasmids and antibodies generated at the MRC PPU at the University of Dundee can be requested through our website (https://mrcppureagents.dundee.ac.uk/). Plasmid request requires a universal material transfer agreement (MTA) that can be completed online at the time of plasmid request. For the purpose of open access, the authors have applied a CC BY 4.0 public copyright license to all Author Accepted Manuscripts arising from this submission.

## Materials and Methods

### Generation of Stable LysoTag Cell Lines

Detailed protocol for preparing the stable cell lines used in this study are described at: dx.doi.org/10.17504/protocols.io.261gedz2ov47/v1 (steps 1 - 22). Briefly: HEK293 cells (The American Type Culture Collection (ATCC). Cat# CRL-1573, RRID:CVCL 0045) were transfected with lentiviral constructs expressing either TMEM192-3×HA (LysoTag) (https://www.addgene.org/102930/) or a control vector expressing only 3×HA (MockTag) (DU70022) and cultured in DMEM containing 10% (by vol) fetal bovine serum supplemented with L-glutamine and penicillin and streptomycin. After 24h cells were subjected to selection with puromycin (2 μg/ml) for ∼72h.

### LysoTagIP method for HEK293 cells and iPSC-derived dopaminergic neurons

We have slightly adapted the previously described LysoTagIP method (10) with the main difference being employing an Isobiotec cell breaker for the homogenization step. Detailed protocol for this slightly modified method is described at: dx.doi.org/10.17504/protocols.io.261gedz2ov47/v1 (Steps 23-52). Briefly: cells stably expressing LysoTag or MockTag (RRID:CVCL_C8A7) were cultured in the continued presence of puromycin (2 μg/ml) in 15 cm diameter dishes to 90% confluency. Cells were scraped into Potassium Phosphate Buffer Saline (KPBS) containing 1X Complete Protease Inhibitor cocktail (Cat#11836170001) and 1X PhosSTOP phosphatase inhibitor cocktail (Cat#4906845001). An aliquot (5% by vol) of the cell suspension was removed and cells pelleted at 1500 x g and frozen in liquid nitrogen and stored at -80 °C for subsequent proteomic, lipidomic or metabolomic analysis. This fraction is termed the “whole cell extract”. The remaining cell suspension was homogenized using Isobiotec cell-breaker. Homogenates were cleared of debris by centrifugation at 1,500 × g for 2 min at 4 °C. The remainder of the supernatant containing cellular organelles was collected and transferred to a clean 1.5 mL Eppendorf tube containing 100 µL of 50% (by vol) slurry of anti-HA magnetic bead (Thermo Fisher Cat# 88837). The mixture was incubated at 4 °C for 5 min on an orbiter ensuring constant gentle agitation. After incubation, the tube containing the mixture was placed on a magnet for 30 s and the supernatant was removed. The beads were washed 3 times with ice cold KPBS, aliquoted and snap frozen in liquid nitrogen and stored in -80 °C until further processing (see below).

The iPSC-derived A53T-α-synuclein dopaminergic neurons and their corresponding isogenic controls (25) were differentiated following the Kriks protocol (26) and as described in recent publications (27, 28). At an age of 84 days after start of the differentiation, cells were harvested and processed for tagless LysoIP as described above.

### Generation of monoclonal TMEM192 antibody-coupled magnetic beads

Detailed protocol for generating monoclonal TMEM192 antibody-coupled magnetic beads is described at dx.doi.org/10.17504/protocols.io.q26g7p2ykgwz/v1. For each coupling reaction, 20 mg of MyOne™ Epoxy Dynabeads™ (Invitrogen™, 34001D) was conjugated with 600 µg of rabbit monoclonal TMEM192 antibody [TMEM192^AB1^ (Abcam ab186737) and/or TMEM192AB2 (Abcam ab185545)]. The antibodies were purchased in PBS and their concentrations adjusted to 1.2 mg/ml. 20 mg of magnetic beads were resuspended in 1 ml of sterile Milli-Q water, vortexed, and sonicated in a water bath sonicator for 5 minutes, after which Mili-Q water was removed. Beads were resuspended in 1 ml of sterile Milli-Q water and the sonication was repeated. After sonication, Mili-Q water was removed, and the beads were mixed with 0.5 ml of 1.2 mg/ml antibody solution (total amount 600 µg of the antibody) and vortexed, after which 500 μl of buffer C2 (3 M Ammonium Sulphate ((NH4)2SO4) in 0.1 M Sodium Phosphate buffer pH 7.4) was added. The beads suspension was vortexed again and then was incubated on a Thermomixer at 37 °C, 1500 rpm for 20 h. After coupling, beads were washed once with buffer HB (100 mM Glycine pH 11.3, 0.01% Tween-20), followed by one wash with buffer LB (200 mM Glycine pH 2.8, 0.01% Tween-20), and 3 washes with buffer SB (50 mM Tris-HCl (NH2C(CH2OH)3·HCl) pH 7.4 with 140 mM NaCl and 0.1% Tween-20). Beads were then resuspended in 2 ml of the storage buffer (50 mM Tris-HCl (NH2C(CH2OH)3·HCl) pH 7.4 with 140 mM NaCl, 0.1% Tween-20 and 0.2% NaN3) to a final concentration of 10 mg/ml, and stored at 4°C. We used the beads stored in this manner for up to one month.

### Generation of BSA-coupled magnetic beads for MockIP controls

The method is identical to the method above describing generation of monoclonal TMEM192 antibody-coupled magnetic beads, except that 600 µg of bovine serum albumin (BSA), buffer exchanged into Phosphate Buffered Saline (PBS) was coupled to 20 mg of MyOne™ Epoxy Dynabeads™ (Invitrogen™, 34001D).

### Tagless LysoIP method for HEK293 cells

The method is identical to the LysoTagIP method described above, except that monoclonal TMEM192 antibody-coupled magnetic beads and BSA-coupled magnetic beads (MockIP) replace the HA-magnetic beads with the same amount and volume of extract and magnetic beads.

### Tagless LysoIP method for peripheral blood mononuclear cells (PBMCs)

Detailed protocols for isolation of PBMCs and tagless LysoIP are described with embedded video method at: dx.doi.org/10.17504/protocols.io.x54v9yp51g3e/v1. Briefly: fresh blood (5-60 ml) was collected from participants into K2EDTA vacutainers and peripheral blood mononuclear cells isolated using density gradient centrifugation as described previously (16) and in protocols.io (29). Isolated cells were briefly washed 3 times with PBS and cells resuspended in 0.8 ml KPBS containing 1X Complete Protease Inhibitor cocktail (Cat#11836170001) and 1X PhosSTOP phosphatase inhibitor cocktail (Cat#4906845001) (volume of resuspension buffer independent of volume of starting blood volume) and supplemented with freshly diluted 0.5 mM diisopropylfluorophosphate made up from 0.5M stock in isopropanol to inhibit elastase proteases derived from low levels of contaminating neutrophils. Note diisopropylfluorophosphate is highly toxic and must be handled in a fume hood and waste disposed in 2% (w/v) NaOH. An aliquot (5% by vol) of the cell suspension was removed and cells pelleted at 1500 x g and frozen in liquid nitrogen and stored at -80 °C for subsequent proteomic, lipidomic or metabolomic analysis. This fraction is termed the “whole cell extract”. The remaining resuspended PBMCs were homogenized using Isobiotec cell-breaker. The remaining steps for the immunoprecipitation are identical to those described above for the tagless LysoIP method. An aliquot (5% by vol) of the homogenate is kept back and termed “whole cell extract” for subsequent proteomic, lipidomic or metabolomic analysis.

### Study participants and blood sample collection

Between 10-60ml of peripheral blood was collected from healthy volunteers for setting up the method. From November 2022 and February 2024, between 8 and 10ml of fresh peripheral blood was collected from a total of 5 participants with CLN3 Batten disease: 2 were recruited from the Department of Metabolic Diseases, Wilhelmina Children’s Hospital, University Medical Center Utrecht, Utrecht University, Utrecht in the Netherlands and 3 from the Department of Pediatrics, University Medical Center Hamburg-Eppendorf, Hamburg, Germany. Additionally, a total 5 sex matched young healthy donors were recruited from both clinical sites as controls (Tab 1).

Diagnosis was confirmed upon identification of bi-allelic homozygous pathogenic variants in the CLN3 gene (CLN3; Chr16(GRCh37)c.461-280_677+382del966). In the case of the patient with adult onset (5th decade) retinal only disease, the diagnosis was based on the combination of bi-allelic compound heterozygous variants in CLN3 (c.391_392del; p.(Ser131fs) and CLN3 (c.969G>A; r. (spl?)), clinical symptoms with characteristic optical coherence tomography (OCT) abnormalities (30, 31) and characteristic fingerprint abnormalities on the electron microscopy of the skin (32). Demographics including sex, age at disease onset, clinical phenotype and CLN3 genotype were collected (Tab. 1).

### Ethical approval and consent to participate

The study was approved by the respective local ethics committees: Medical ethics committees of the Ärztekammer Hamburg, Germany (PV7215) and the NedMec, to which the UMC Utrecht is affiliated (METC, 23-268/A). Patients’ or parentś written informed consent was obtained according to the Declaration of Helsinki (1991). Additional, non-clinical research ethics approval was in place for experiments using blood from healthy volunteers to set up the methodology (University of Dundee, SMED REC Number 22/84).

**Flow cytometry assay**

Detailed protocol for the flow cytometry assay is described at dx.doi.org/10.17504/protocols.io.n2bvj378xlk5/v1. HEK293 cells (15 cm dish, 90% confluent) and PBMCs (isolated from 15 ml of serum and divided into 4 equal aliquots) were treated ± 200 nM Bafilomycin at 37 C° for 3.5 h and cells then incubated for a further 30 min ± 50 nM Deep Red LysoTracker at 37 C° for 30 min. Both bafilomycin and LysoTracker were dissolved in DMSO at a 1000X concentration and for the minus-bafilomycin and/or minus-Lysotracker conditions the equivalent volume of DMSO was added. Tagless LysoIP were performed as described above. The isolated lysosomes bound to the magnetics beads were diluted 1 in 10 in KPBS and transferred to FACS tubes and analyzed on a BD LSR Fortessa cytometer at 647 nm excitation measuring the emission at 668 nm, the optimal wavelength for Deep Red LysoTracker.

### Cathepsin D activity assay

Detailed protocol for the Cathepsin D assay is described at: dx.doi.org/10.17504/protocols.io.8epv5r7j5g1b/v1. Wildtype HEK293 cells (15 cm dish, 90% confluent) were subjected to tagless LysoIP as described above. The isolated lysosomes bound to the magnetics beads were diluted 1 in 10 in KPBS. Protein levels in the whole cell extracts and LysoIP were determined using the ultrasensitive Bicinchoninic acid assay (BCA) method (33). 2 µg of protein from the whole cell extract and the LysoIP, MockIP and WCL were aliquoted into wells in a 96-well plate and assayed using a fluorometric Cathepsin D activity assay kit (Abcam. Cat# Ab65302) that is based on quantifying the hydrolysis of the preferred cathepsin-D substrate sequence GKPILFFRLK(Dnp)-D-R-NH2) labeled with MCA fluorescent dye. Cathepsin D substrate hydrolysis released fluorophore fluorescence was measured on a Clariostar Plate Reader (Ex/Em= 328 nm/460 nm).

### Glucocerebrosidase-1 enzyme activity assay

Detailed protocol for the glucocerebrosidase-1 enzyme activity assay is described at: https://dx.doi.org/10.17504/protocols.io.8epv5r9jdg1b/v1. Glucocerebrosidase-1 (GCase) enzyme activity was measured by monitoring the cleavage of the fluorescent substrate 4-methylumbelliferyl-β-D-glucopyranoside (4-MUG). Detailed protocol for the GCase enzyme activity assay is described at (34). Briefly, PBMCs were isolated from 15 ml fresh peripheral blood from 6 healthy volunteers and subjected to tagless LysoIP as described above. The isolated lysosomes and the whole cell pellets were suspended in 1% (v/v) Triton X-100 lysis buffer, cleared by centrifugation, and protein concentrations were determined using the ultrasensitive BCA method (33). 5 µg of protein from the whole cell extract and 1 µg of protein from the LysoIPs were aliquoted into a 96-well plate in duplicates, and incubated with 500 µM 4-MUG, in assay buffer (0.15 M citrate-phosphate buffer, pH 5.4, 0.25% (w/v) sodium taurocholate, 1 mM EDTA, 0.5% (w/v) BSA), in the presence or absence of 300 µM conduritol β-epoxide (CBE, glucocerebrosidase-1 inhibitor), at 37 °C. After 1h incubation, the reaction was stopped by addition of 1 M glycine, pH 12.5, calibrators [0 – 10 µM 4-methylumbelliferone (4-MU)] were aliquoted into empty wells of the plate, and the fluorescence intensity was measured on a Pherastar plate reader (Ex/Em = 350/460). GCase activity was estimated from the generated calibration curve as the amount of released fluorophore (4-MU) per mg of protein extract per min of reaction.

### Immunofluorescence assay

Detailed protocol for the immunofluorescence assay is described at: dx.doi.org/10.17504/protocols.io.q26g71zykgwz/v1. Briefly, HEK293 cells (ATCC. Cat# CRL-1573, RRID:CVCL_0045) were grown on Poly-L-lysine coated 22 × 22 glass coverslips in 6-well 3.5 cm diameter plates. For fixation, medium was aspirated, and cells were fixed in 3 ml of 4% (w/v) paraformaldehyde in PBS for 10 minutes at room temperature. Fixed cells were then washed three times at 5 minutes intervals with 0.2% (w/v) bovine serum albumin (BSA) dissolved in PBS. Cells were permeabilized with 1% (v/v) Nonidet P40 diluted in PBS for 10 minutes at room temperature. Permeabilized cells were blocked with 1% (w/v) BSA in PBS for 1 h, then incubated with primary antibody at 1:1000 dilution in PBS for 1 h at room temperature in a dark chamber. The combination of the primary antibodies used is Mouse anti-LAMP1 (Santa Cruz. Cat#sc-20011. RRID:AB_626853) and Rabbit anti-TMEM192 (Abcam. Cat# Ab186737. RRID:AB_3095637) or Mouse anti-ATPB (Abcam. Cat# Ab14730. RRID:AB_301438) and Rabbit anti-TMEM192 (Abcam. Cat# Ab186737. RRID:AB_3095637). This is followed by three washes with 0.2% (w/v) BSA at 5 minutes intervals. Cells were then incubated for 1 h, in a dark chamber, with a mixture of secondary antibodies containing Alexa Fluor 488 donkey anti-mouse (Invitrogen. Cat# A21206. RRID:AB_2535792) and Alexa Fluor 594 goat anti-rabbit (Invitrogen Cat# A11012. RRID:AB_2534079) at 1:500 dilution and Hoechst 33342 (Thermo Fisher. Cat# 62249) at 1:1000 in PBS. Cells were washed again three times with 0.2% (w/v) BSA, rinsed in MilliQ water, and mounted on glass microscope slides with VECTASHIELD antifade mounting media (Vector Laboratories, H1000). Slides were then imaged using Leica TCS SP8 MP Multiphoton Microscope using a 40x oil immersion lens choosing the optimal imaging resolution with 1-pixel size of 63.3 nm × 63.3 nm.

### Sample preparation and analysis for quantitative proteomics

Detailed protocol for the processing of the whole cell extract, LysoTagIP and LysoIP is described at: dx.doi.org/10.17504/protocols.io.q26g7p2d8gwz/v1. A detailed protocol for the data independent mass spectrometry analysis is described at: dx.doi.org/10.17504/protocols.io.kxygxzrokv8j/v1. The stored and frozen whole cell extract pellets and the LysoTagIP, LysoIP and MockIP samples from wildtype and LysoTag HEK293 cells, PBMCs were resuspended in 0.1ml of 2% (w/v) (sodium dodecyl sulfate) SDS, 20 mM HEPES pH 8.0 containing 1X Complete Protease Inhibitor cocktail (Cat#11836170001) and 1X PhosSTOP phosphatase inhibitor cocktail (Cat#4906845001). Tryptic digestion was carried out using S-Trap-assisted On-column tryptic digestion (35, 36). The digested tryptic peptides were then analyzed on Orbitrap Exploris 480, tims-TOF Pro, tims-TOF SCP and Orbitrap Astral mass spectrometers. The details of LC column, DIA isolation window acquisition schemes and mass spectrometer data acquisition parameters are provided in supplemental file MSSettings. The raw MS data were further processed using the DIA-NN (1.8.1) (37) against the Uniprot Human database (downloaded January 2023; 20381 entries with isoforms) in a library free mode. DIA-NN database search parameters for each dataset are provided in supplemental file MSSettings. The output files from DIA-NN search were further processed using Perseus software suite (version 1.6.15.0) (38) for differential analysis two-sided t-Test (Permutation FDR = 0.01,S0=0.1), lysosomal annotated proteins were further Z-score normalized and subsequently used for supervised heatmap clustering and violin plot representation of LysoIP, LysoTag-IP and WCLs. Further, data visualization was performed using CURTAIN tool (39) and Graphpad prism software suites.

### Sample preparation and analysis for quantitative lipidomics

Detailed protocol for the processing of the whole cell extract, LysoTagIP and LysoIP is described at: dx.doi.org/10.17504/protocols.io.yxmvm3r8bl3p/v1. The stored and frozen whole cell extract pellets and the LysoTagIP, LysoIP and MockIP samples were resuspended in 1 ml LC-MS grade chloroform/methanol (2:1 by volume) supplemented with 1 μg/ml Splashmix internal standard (Avanti, #330707-1EA). Lipids were profiled using an Acentis Express C18 150 x 2.1 m column (Millipore Sigma 53825-U) and the ID-X Tribrid mass spectrometer. Unbiased differential analysis was performed by LipidSearch and Compound Discoverer. Normalization was performed by constant median after blank exclusion. Targeted BMP analyses were performed from an adapted protocol (40) using an Agilent C18 column (Agilent Technologies 821725-90) and Ultivo triple quadrupole mass spectrometer.

### Sample preparation and analysis for quantitative metabolomics

Detailed protocol for the processing of the whole cell extract, LysoTagIP and LysoIP is described at: dx.doi.org/10.17504/protocols.io.kxygx3jm4g8j/v1. The stored and frozen whole cell extract pellets and the LysoTagIP, LysoIP and MockIP samples were resuspended in LC-MS grade 80% methanol (v/v) with isotopically labeled amino acids (Cambridge Isotope Laboratories, Inc. Catalog #MSK-A2-1.2). Metabolites were profiled by liquid chromatography/mass spectrometry (LC/MS) using a SeQuant® ZIC®-pHILIC 150 x 2.1 mm column (Millipore Sigma 1504600001) and an ID-X Tribrid mass spectrometer. Unbiased differential analysis was performed in Compound Discoverer (ThermoFisher Scientific). Rigorous quantification of metabolite abundance was performed by TraceFinder.

### Data visualization, statistics and figure generation

Perseus 1.6.15.0 version was used for proteomics data analysis and CURTAIN tool was used for proteomic data visualization. GraphPad Prism version 10.0.0 for macOS, GraphPad Software, Boston, Massachusetts USA, www.graphpad.com was used for data visualization and statistics. The type of analysis is provided in each figure legend. Workflow figures and schematics were created using BioRender.com.

## Supplementary Figures

**Fig. S1.**
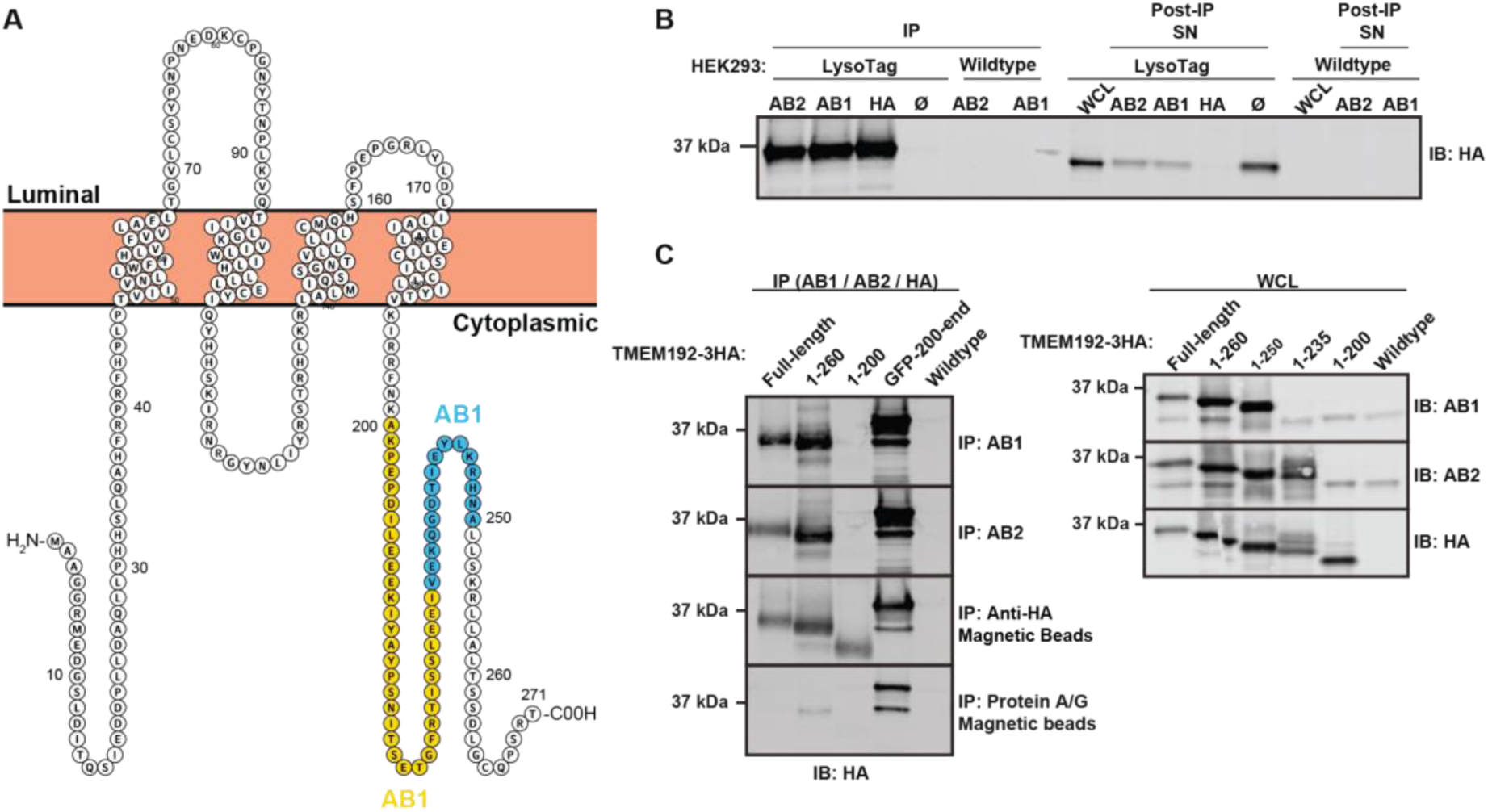
Characterization of 2 commercially available TMEM192 antibodies for their ability to immunoprecipitate TMEM192. **(A)** Domain architecture of TMEM192 protein with regions containing antibody epitopes highlighted in blue (AB1, Abcam ab186737) and yellow (AB2, Abcam ab185545) (Figure created using Protter wlab.ethz.ch/protter). **(B)** Ability of the two TMEM192 antibodies to immunoprecipitate overexpressed TMEM192. TMEM192-3HA was immunoprecipitated from HEK293 lysates using AB1, AB2 and HA beads, using empty beads (Ø) and HEK293 lysates without overexpression (Wildtype) as negative controls. Immunodepletion was additionally assessed in the post-IP supernatants (SN). **(C)** Epitope analysis to map the respective TMEM192 antibody binding sites. Full-length and the indicated C-terminal truncations of TMEM192-3HA were overexpressed and immunoprecipitated using AB1, AB2 and HA beads to confirm the presence of the epitopes within the C-terminal region of TMEM192. Further analysis was performed by immunoblotting of a panel of overexpressed truncations (1-260, 1-250, 1-235, 1-200) and full-length TMEM192-3HA using AB1, AB2 and HA antibodies.

**Fig. S2.**
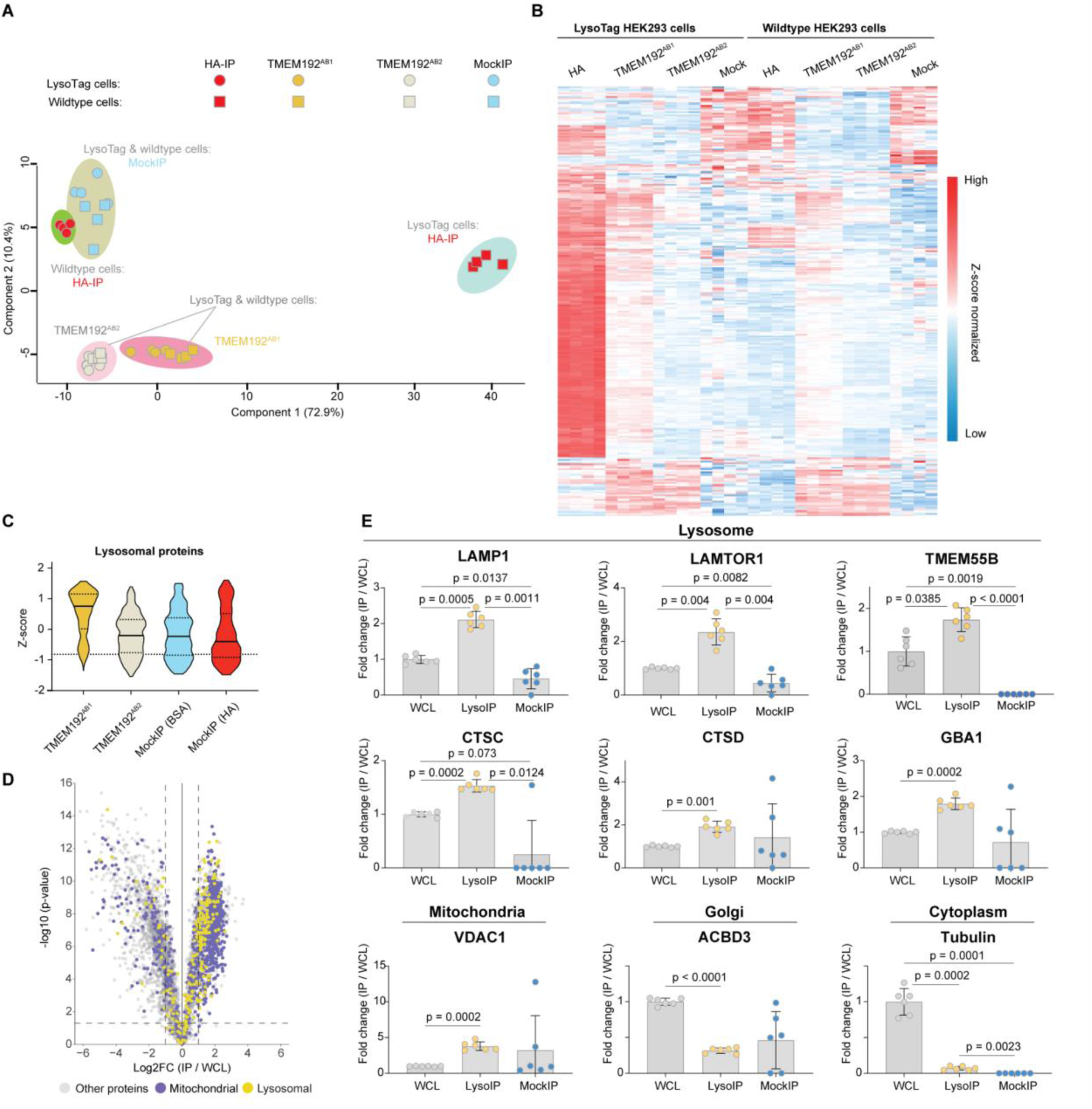
TMEM192 antibody selection for tagless LysoIP. **(A)** Principal component analysis of DIA mass spectrometry data of immunoprecipitates using TMEM192^AB1^/ TMEM192^AB2^/ HA-/ BSA-coupled magnetic beads in wildtype HEK293 cells (filled squares n = 4) and LysoTag HEK293 cells (filled circles n = 4). **(B)** Heatmap clustering of lysosomal annotated proteins denoting significant enrichment of lysosomal proteins in HA-IP and TMEM192^AB1^ IP (Z-score normalized). **(C)** Similar to B, the violin plot depicting the enrichment of curated lysosomal-annotated proteins described in Dataset S1 using the different coupled magnetic beads in wildtype HEK293 cells (n = 4). **(D)** Volcano plot showing the fold enrichment/depletion of proteins in the LysoIPs (n = 6, p-value adjusted for 1% permutation-based FDR correction, s0 = 0.1). The yellow dots indicate known lysosomal-annotated proteins, and the purple dots indicate mitochondrial-annotated proteins curated from databases described in Datasets S1 and S2 (https://doi.org/10.5281/zenodo.11085342). Curtain link: https://curtain.proteo.info/#/f097ad91-ed49-4a29-bb4b-ed8037009a04. **(E)** Bar charts depicting the relative enrichment of indicated protein markers for lysosome, mitochondria, Golgi and cytoplasm. Data presented as mean ± SEM (n = 6). One-way ANOVA with Tukey’s HSD post-hoc was used for multiple comparison analysis between the groups.

**Fig. S3.**
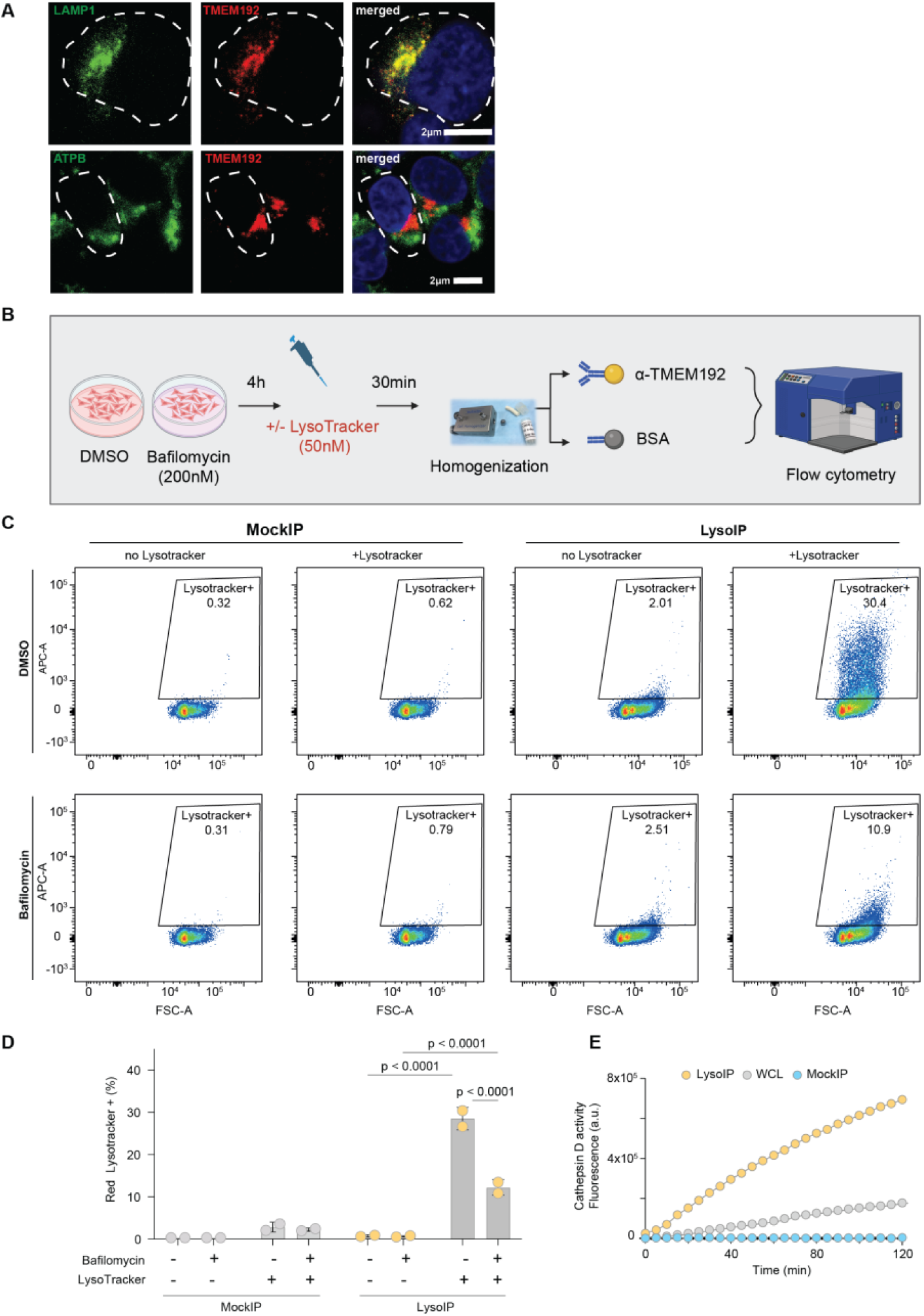
Isolation of functional lysosomes using the endogenous TMEM192-IP in HEK293 cells. **(A)** Anti-TMEM192^AB1^ is specific for the lysosome and not mitochondria. Scale bar = 2 μm. **(B)** Schematic of workflow for flow cytometry analysis in HEK293 cells. **(C)** Quantitation of the percentage of beads positive for the LysoTracker. **(D)** Representative scatter plot from one experimental replicate (n = 2). Ordinary two-way ANOVA with Tukey’s multiple comparison test. E) Measurement of Cathepsin D activity in LysoIP, MockIP and WCLs of HEK293 cells (n = 2).

**Fig. S4.**
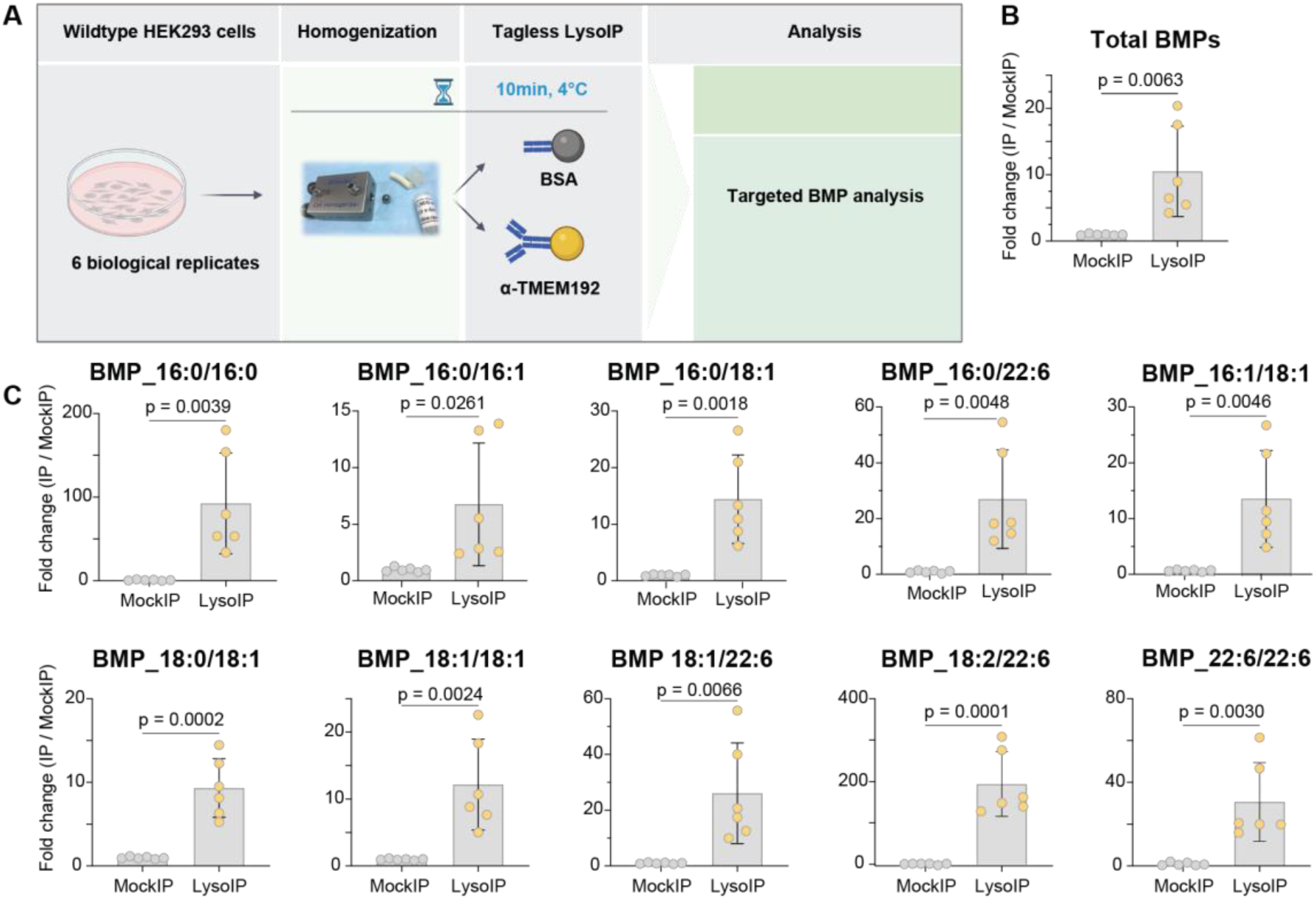
Bis(monoacylglycero)phosphates are enriched in LysoIP samples from wildtype HEK293 cells. **(A)** Experimental design to study lysosomal lipids using tagless LysoIP followed by targeted BMP analysis. **(B)** Targeted analysis of accumulated BMPs in lysosomes derived from wildtype HEK293 cells using tagless LysoIP (n = 6). Relative enrichment of total BMPs quantified in the LysoIP samples and the MockIP samples. Data presented as mean ± SEM (n = 6). Statistical analysis was performed using Student’s t-test. **(C)** Enrichment of specific BMP species in the LysoIP samples.

**Fig. S5.**
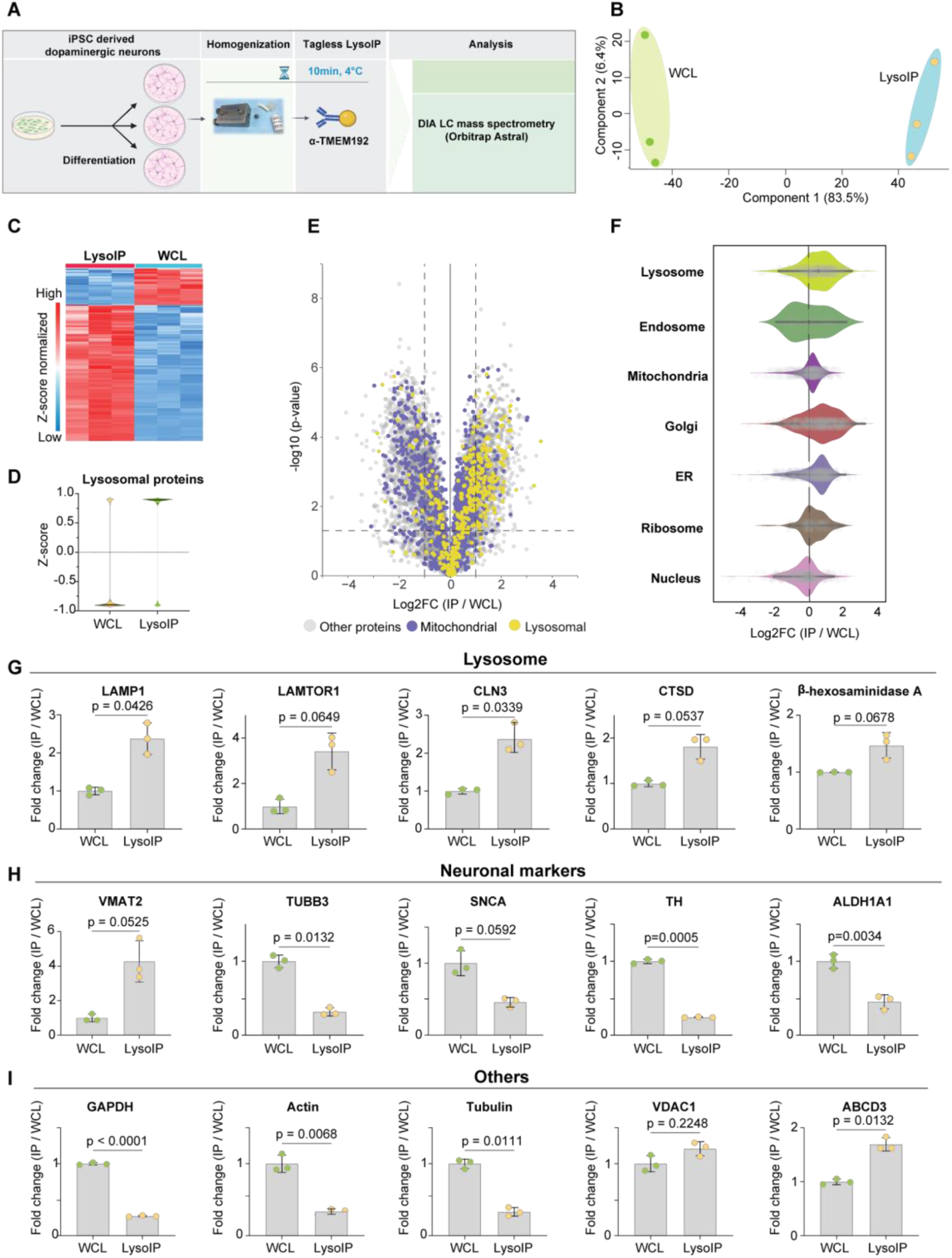
Tagless LysoIP from human iPSC-derived dopaminergic neurons. **(A)** Workflow. **(B)** Principal component analysis of DIA mass-spectrometry data of LysoIPs from iPSC-derived dopaminergic neurons (n = 3). **(C)** Volcano plot of proteins enriched/depleted in the LysoIPs compared to whole cell extracts (WCL). Curtain link: https://curtain.proteo.info/#/c4ee2603-109a-4a7a-b762-247735b38ab0. **(D)** Protein profile heatmap. **(E)** Violin plots of lysosomal proteins enriched via the tagless LysoIP in the immunoprecipitates from iPSC-derived dopaminergic neurons. **(F)** Organelle profiling demonstrates enrichment of lysosomal proteins and modest enrichment of other organelles. Bar plots of representative **(G)** lysosomal proteins, **(H)** neuronal markers and **(I)** cytosolic, mitochondrial and Golgi proteins in LysoIPs and their respective whole cell extracts (WCL). Data presented as mean ± SEM (n = 3). Statistical analysis was performed using Student’s t-test.

**Fig. S6.**
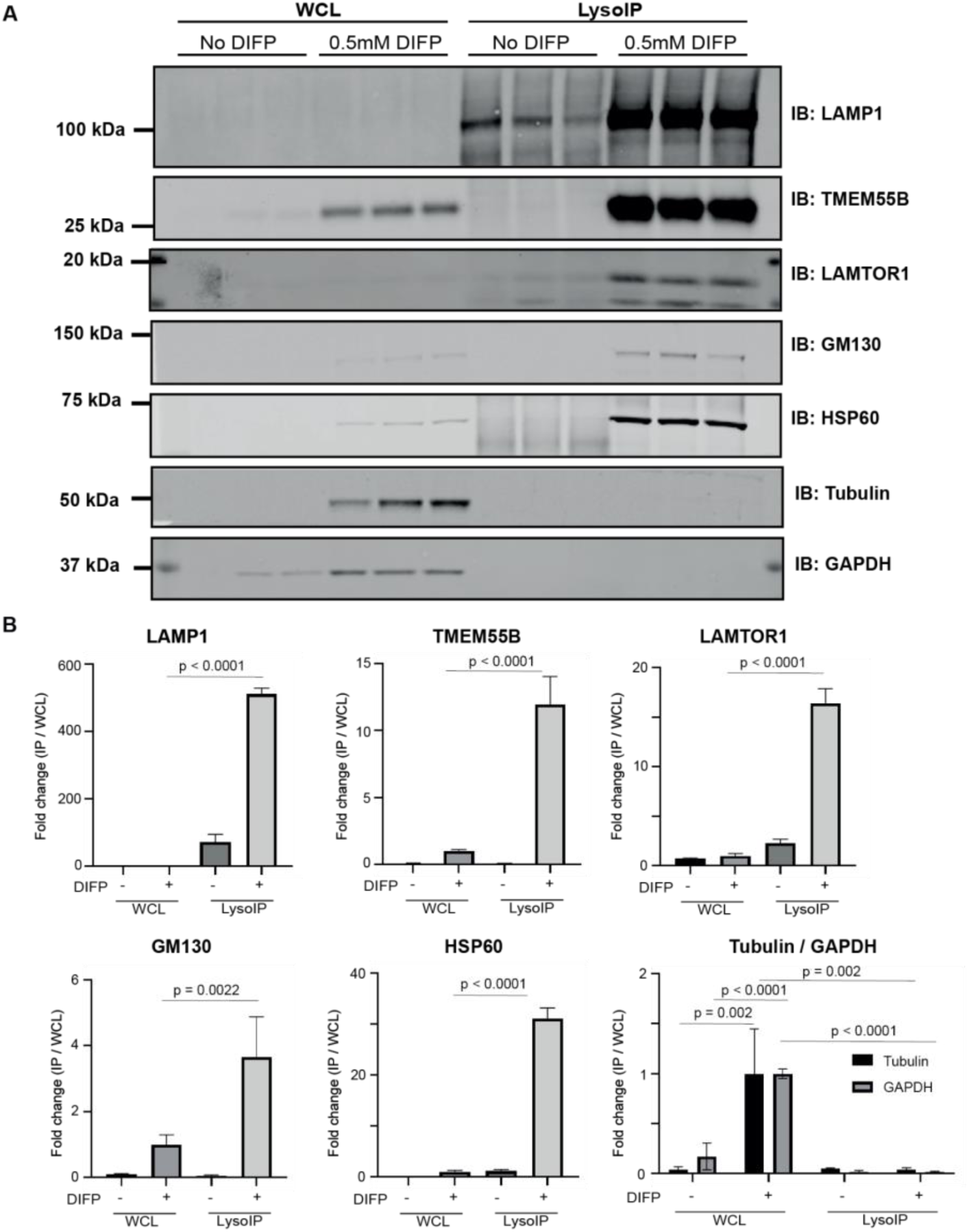
Diisopropylfluorophosphate is required to prevent protein degradation in PBMCs. **(A)** Immunoblot analysis confirming enrichment of lysosomes in the LysoIP in PBMCs. PBMCs were treated with 0.5 mM diisopropylfluorophosphate (DIFP) to prevent rapid degradation of proteins. Whole-cell lysates (2 µg) as well as the resuspended immunoprecipitates (IPs) (2 µg) were subjected to immunoblotting with the lysosomal (LAMP1, TMEM55B, LAMTOR1), Golgi (GM130), cytosolic (α-tubulin, GAPDH) and mitochondrial (HSP60) markers. The data shown is from 3 healthy male donors. **(B)** Quantitative immunoblotting analysis. The graph represents ratios of IP/whole-cell lysate (mean ± SEM, n = 3). One-way ANOVA with Tukey’s HSD post-hoc was used for multiple comparison analysis between the groups.

**Fig. S7.**
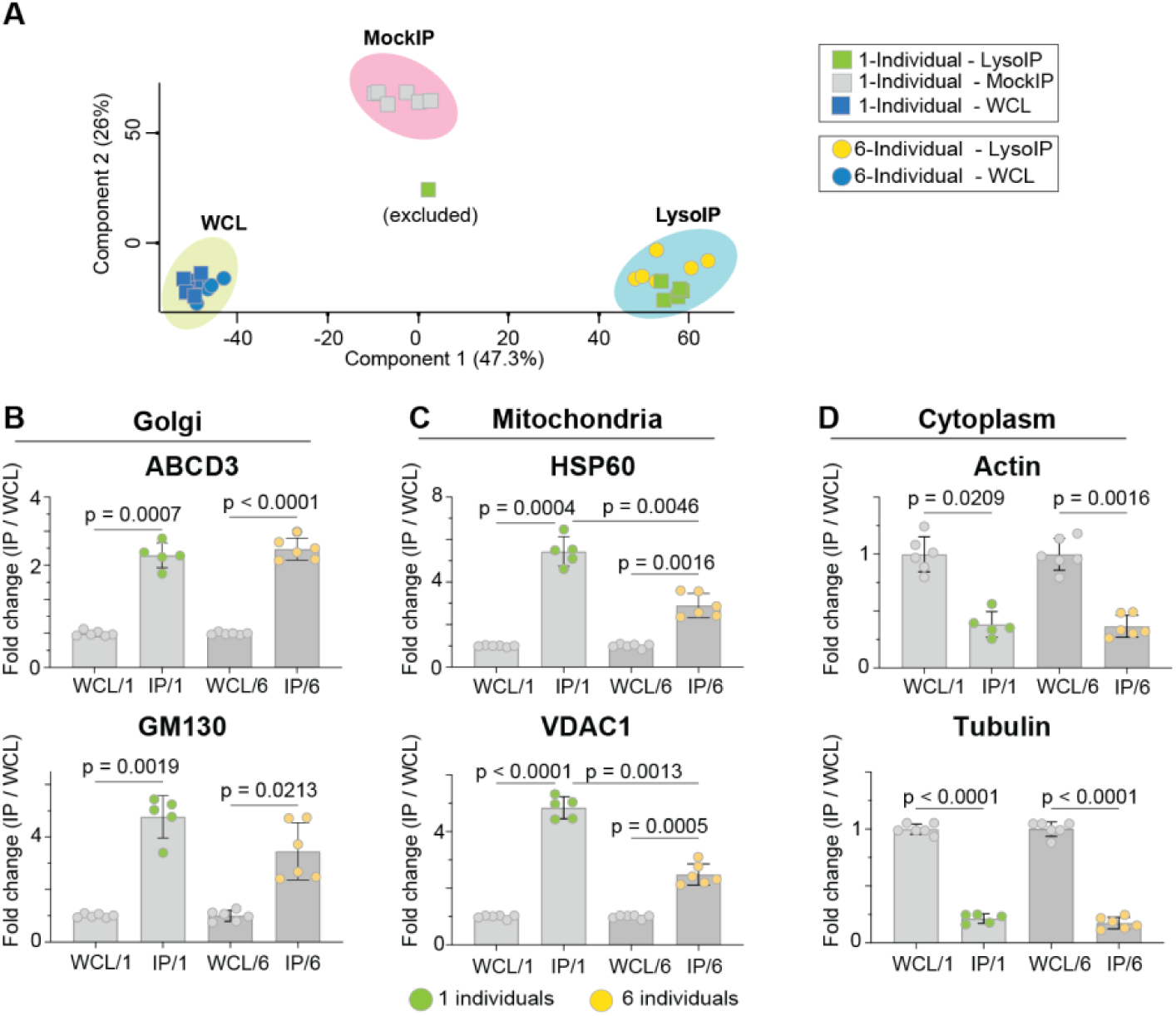
Characterization of the tagless LysoIP in PBMCs from healthy donors: **(A)** Principal component analysis of DIA mass spectrometry data of LysoIP, MockIP (for single donor experiment only) immunoprecipitates as well as WCLs. **(B)** Bar graphs of representative proteins from Golgi-, mitochondria and cytosol enriched/depleted in the LysoIPs. The graph represents ratios of IP/WCL (mean ± SEM, n = 6). One-way ANOVA with Tukey’s HSD post-hoc was used for multiple comparison analysis between the groups.

